# Effects of age on responses of principal cells of the mouse anteroventral cochlear nucleus in quiet and noise

**DOI:** 10.1101/2024.05.22.595362

**Authors:** Maggie Postolache, Catherine J. Connelly Graham, Kali Burke, Amanda M. Lauer, Matthew A. Xu-Friedman

## Abstract

Older listeners often report difficulties understanding speech in noisy environments. It is important to identify where in the auditory pathway hearing-in-noise deficits arise to develop appropriate therapies. We tested how encoding of sounds is affected by masking noise at early stages of the auditory pathway by recording responses of principal cells in the anteroventral cochlear nucleus (AVCN) of aging CBA/CaJ and C57BL/6J mice *in vivo*. Previous work indicated that masking noise shifts the dynamic range of single auditory nerve fibers (ANFs), leading to elevated tone thresholds. We hypothesized that such threshold shifts could contribute to increased hearing-in-noise deficits with age if susceptibility to masking increased in AVCN units. We tested this by recording the responses of AVCN principal neurons to tones in the presence and absence of masking noise. Surprisingly, we found that masker-induced threshold shifts decreased with age in primary-like units and did not change in choppers. In addition, spontaneous activity decreased in primary-like and chopper units of old mice, with no change in dynamic range or tuning precision. In C57 mice, which undergo early onset hearing loss, units showed similar changes in threshold and spontaneous rate at younger ages, suggesting they were related to hearing loss and not simply aging. These findings suggest that sound information carried by AVCN principal cells remains largely unchanged with age. Therefore, hearing-in-noise deficits may result from other changes during aging, such as distorted across-channel input from the cochlea and changes in sound coding at later stages of the auditory pathway.

**Significance Statement:** Middle age and older listeners commonly experience hearing deficits in the presence of background noise. Central auditory areas have been implicated in hearing-in-noise deficits, but it is not known where these deficits arise. We performed *in vivo* recordings in mice of different ages at the first stage of the auditory pathway in the brain, the cochlear nucleus, to examine how encoding of sounds is perturbed by masking noise. We found that the responses of individual neurons remain largely intact with age, including the processing of tones in masking noise, despite previously documented structural and physiological degeneration of their auditory nerve inputs. This suggests that problems hearing in masking noise result from changes at other stages of the auditory pathway.

## Introduction

Age-related decline of the auditory system affects millions of people worldwide. Age-related changes include deficits in temporal processing (Strouse et al., 1998; Walton et al., 1998; Frisina and Walton, 2006; Anderson and Karawani, 2020) and poor hearing in noisy conditions (Kobrina et al., 2020; Shilling-Scrivo et al., 2021, 2022). These declines may result from changes in both the cochlea and central auditory pathway. Age-dependent changes in the cochlea are well-documented, including loss of inner and outer hair cells (Wu et al., 2019; Wang and Puel, 2020). In addition, inner hair cells contact fewer auditory nerve fibers (ANFs) (Bao and Ohlemiller, 2010; Sergeyenko et al., 2013; Kobrina et al., 2020). The spontaneous activity and sensitivity of ANFs decrease in aged gerbils, but for sounds above threshold, temporal coding and tuning appear similar to younger animals (Heeringa et al., 2020; Heeringa et al., 2023). Thus, age-dependent hearing loss is not fully accounted for by cochlear degeneration, and it is important to determine where deficits occur to identify treatments to improve hearing in older listeners.

Hearing speech in masking noise requires accurate processing of the timing, intensity, spectral content, and location of sounds. Processing of these features depends critically on two targets of ANFs in the anteroventral cochlear nucleus (AVCN), bushy and stellate cells (Young and Oertel, 2004), which correspond to primary-like and chopper units in *in vivo* recordings (Blackburn and Sachs, 1989). Bushy cells convey precise temporal information to the superior olive for sound localization (Oertel, 1999). Stellate cells encode the envelopes of sounds important for speech perception (Wang and Sachs, 1994; Shannon et al., 1995; Oertel et al., 2011) and contribute to the olivocochlear efferent feedback reflex (Warr, 1972; Darrow et al., 2012; Romero and Trussell, 2022). Bushy and stellate cells integrate excitation from multiple ANFs with inhibition from a variety of sources, yielding distinct spectral and temporal responses (Wickesberg and Oertel, 1990; Kopp-Scheinpflug et al., 2002; Dehmel et al., 2010; Xie and Manis, 2013; Keine and Rubsamen, 2015). Inhibition throughout the auditory pathway appears to change with age (Caspary et al., 2008), which could cause changes in AVCN activity that could contribute to deficits in auditory processing in older listeners.

Masking noise affects responses of ANFs and cochlear nucleus units to tones, and shifts their dynamic ranges to higher intensity levels (Costalupes et al., 1984; Ehret and Moffat, 1984; Gibson et al., 1985; Young and Barta, 1986; Huet et al., 2018). It is unknown how the effects of masking noise change with age. If thresholds shift even more in older animals, it could contribute to deficits hearing in noise.

There is some evidence that cochlear nucleus activity changes with age. Auditory brainstem recordings show smaller and more delayed peaks associated with the cochlear nucleus (Ng et al., 2015). Spontaneous activity appears to change with age at multiple stages of the auditory pathway, decreasing in the auditory nerve in gerbils (Heeringa and Koppl, 2019), and increasing in auditory cortex (Overton and Recanzone, 2016), which could change the balance of excitation and inhibition in complex ways. At the cellular level, the synapses formed by ANFs onto bushy cells, called endbulbs of Held, show atrophy that coincides with the progression of hearing loss (Connelly et al., 2017), while the overall density and number of excitatory and inhibitory synapses appear relatively similar (Helfert et al., 2003; Muniak et al., 2018). Endbulbs in older mice produce smaller EPSPs (Wang et al., 2021) and with somewhat reduced temporal precision (Xie, 2016; Wang et al., 2019). Such structural and functional changes could affect susceptibility to masking noise in older animals.

To address this, we assessed the effects of masking noise on the responses of AVCN principal neurons in young and old mice. We used CBA/CaJ mice, which show gradual hearing loss (Kobrina and Dent, 2020; Kobrina et al., 2021), as well as C57BL/6J (C57) mice, that show early-onset hearing loss (Noben-Trauth et al., 2003; Prosen et al., 2003; Noben-Trauth and Johnson, 2009). We hypothesized that hearing in noise would worsen with age through increased shifts in threshold in both strains. However, we found instead that threshold shifts in masking noise changed little or even decreased with age. In addition, spontaneous activity in AVCN principal neurons changed little or decreased, while dynamic range and tuning precision changed little. These findings suggest that principal neurons of the AVCN encode stimuli in noisy backgrounds similarly in old and young animals. Therefore, age-related hearing-in-noise deficits likely originate elsewhere.

## Methods

### Subjects

We used CBA/CaJ and C57 mice of either sex. To ensure balanced groups, mouse cages were designated for the experimenter that matched the age range, sex, and strain required for experiments. The experimenter randomly selected mice from those cages. All mice were housed in the same holding room and exposed to similar noise levels from daily room checks and weekly cage changes. Mice were housed in groups of 2-3 per cage and provided with food and water *ad libitum*. For the CBA/CaJ strain, experimental age groups were 3–6 months (20 mice), 12–14 months (“1 y”, 13 mice), and 18–20 months (“1.5 y”, 12 mice). For the C57 strain, experimental age groups were 3–4 months (14 mice) and 8–9 months (7 mice). All experiments were approved by the Institutional Animal Care and Use Committee at the University at Buffalo.

### Surgery

Mice were anesthetized intraperitoneally with 200 mg/kg ketamine (KetaVed, Vedco, Inc.) and 10 mg/kg xylazine (AnaSed, Akron Animal Health) and placed on a platform that was kept at 37°C using a circulating water bath (Gaymar T/Pump). Ketamine/xylazine boosters were injected intramuscularly to maintain sedation following positive paw reflexes. Subcutaneous isotonic saline injections (0.3 to 0.5 ml) were administered every hour. The head was secured using jaw cuffs and bite bars (Stoelting Co.). A lateral incision was made in the scalp to expose the skull. A hole was drilled in the skull directly above the left cochlear nucleus (6.0 mm posterior and 2.25 mm lateral of bregma), and the dura mater was removed. Mice were euthanized after each recording session.

### In vivo recordings

Sharp electrodes were prepared on a Sutter P-97 puller using borosilicate glass (OD 1.5 mm, ID 0.86 mm, with filament) and filled with 3 M KCl, with resistances of 5 to 15 MΩ. The electrode was mounted on a Multiclamp 700B headstage attached to a Scientifica IVM on a Patchstar motorized manipulator. The electrode was lowered into the brain to a depth of approximately 4 mm. Units were isolated using a brief noise pulse as a search stimulus. Primary-like and chopper units were identified using typical response characteristics (Blackburn and Sachs, 1989), and recorded in the same area and at a similar depth in the AVCN. We examined spike amplitudes and firing patterns in response to sound to ensure recordings were made from single units.

Tone stimuli were generated through custom-written software running in Igor (Wavemetrics) controlling a PCI-6229 data acquisition system (National Instruments). Tone stimuli were amplified by a Crown XLS1000 amplifier (Crown Audio) and delivered through a Fostex FT28D speaker. Tone intensities were calibrated with a Larson Davis 820 sound level meter. The tone speaker was positioned 23 cm from the left ear. Evoked responses of units were recorded for tones of 2.5 to 24 kHz (0.5 kHz steps), with intensity steps of 5 dB to derive tuning curves, characteristic frequency (CF), and tuning precision (*Q*_10dB_, i.e. the CF divided by the frequency range yielding significant responses at 10 dB above threshold).

Masking noise was generated using a custom noise box, which read an input voltage of 0 to 10.24 V and rapidly generated random voltages scaled by the input. This allowed us to specify noise intensity with high speed and resolution. The device was based on an Arduino Due controlling Texas Instruments chips ADS8681 and DAC8830, and showed rapid cycle times (18 µs, 55 kHz). The noise box output was amplified by the second channel of the Crown amplifier and delivered through a second Fostex FT28D speaker. We assessed the frequency content of the noise between 1 and 20 kHz using a Larson-Davis 820 sound level meter, and the output was within ±5 dB over that range. We determined each unit’s threshold response to noise bursts, and presented masking noise at 20 or 30 dB above threshold. We played CF tones at 5 dB steps to generate a rate-level function, alternating intensity sweeps with and without masking noise. The baseline firing rate was measured after each intensity sweep for 1 s, typically for a total of ∼30 s. Threshold was determined as the minimum tone intensity with firing rate significantly above baseline. Threshold shift was the difference between thresholds measured with and without masking noise. We quantified dynamic range by fitting the rate-level function to a sigmoid curve and assessing the difference in tone intensities associated with firing rates at 10% and 90% of the maximum driven rate. Dynamic range was measured only in units showing saturation at the highest intensities.

Cochlear health was monitored throughout experiments using distortion product otoacoustic emissions (DPOAEs). For most experiments, we used f_1_ = 13.3 kHz at 90 dB SPL, and f_2_ = 15.96 kHz at 80 dB SPL. For experiments with 8 mo. C57 mice, we used f_1_ = 7 kHz and f_2_ = 8.4 kHz, to avoid high-frequency hearing loss common in the strain. Tones were generated by a USB 6361 (National Instruments) and delivered by an ER-10C DPOAE probe driver-preamp (Etymotic Research Inc.). DPOAEs were detected by the integrated microphone and digitized. Experiments were terminated if responses at the difference tone (2f_1_ – f_2_) were not visible.

### Statistical analyses

Significant elevations in spike rate over baseline were determined using a Poisson conditional test (Krishnamoorthy and Thomson, 2004) and confirmed by examining raster plots at intensities near threshold. Normality of data was evaluated using the Shapiro-Wilks test. Most data were not normally distributed, so non-parametric tests were used for most comparisons, including threshold in quiet, threshold shift, spontaneous rate, dynamic range, tuning precision, and first spike latency and jitter. For comparisons between the three age groups of CBA/CaJ mice, we used a Kruskal-Wallis (K-W) test followed by Dunn’s post-hoc test. For comparisons between the two age groups of C57 mice, we used Mann-Whitney (M-W) U tests. Significance of threshold shifts (Figs 3B, 4B) was evaluated using paired Wilcoxon signed rank tests. Summary plots show individual data, with median and median absolute deviation. In some cases, we used linear regression to determine how two variables were related. Statistical values were calculated using SPSS (IBM), with α = 0.05.

## Results

### Characterizing AVCN units

We first investigated how responses of units in the AVCN are affected by age. We performed single-unit extracellular recordings and classified units based on spike shape and response kinetics. Primary-like units are characterized by a spike waveform with a small potential that precedes it (Fig. 1Ai), which is thought to be an extracellularly-recorded EPSP (Kopp-Scheinpflug et al., 2002; Typlt et al., 2010), while chopper units have a simple, biphasic spike (Fig. 1Bi) (Pfeiffer, 1966). We assessed each unit’s tuning curve to identify its characteristic frequency (CF), i.e. the frequency the unit was most sensitive to (Fig. 1Aii, Bii). Figure 1Aii shows the tuning curve from a representative primary-like unit from a 3–6 mo. CBA/CaJ mouse, which had CF of 22 kHz and a threshold of –11.6 dB SPL. Figure 1Bii shows a representative tuning curve from a chopper unit from a 1.5 y CBA/CaJ mouse, which had a CF of 15.5 kHz with a threshold of 37.2 dB SPL. We also used the tuning curve to assess tuning precision (*Q*_10dB_, double-headed arrows in Fig. 1Aii, Bii).

**Figure 1.**
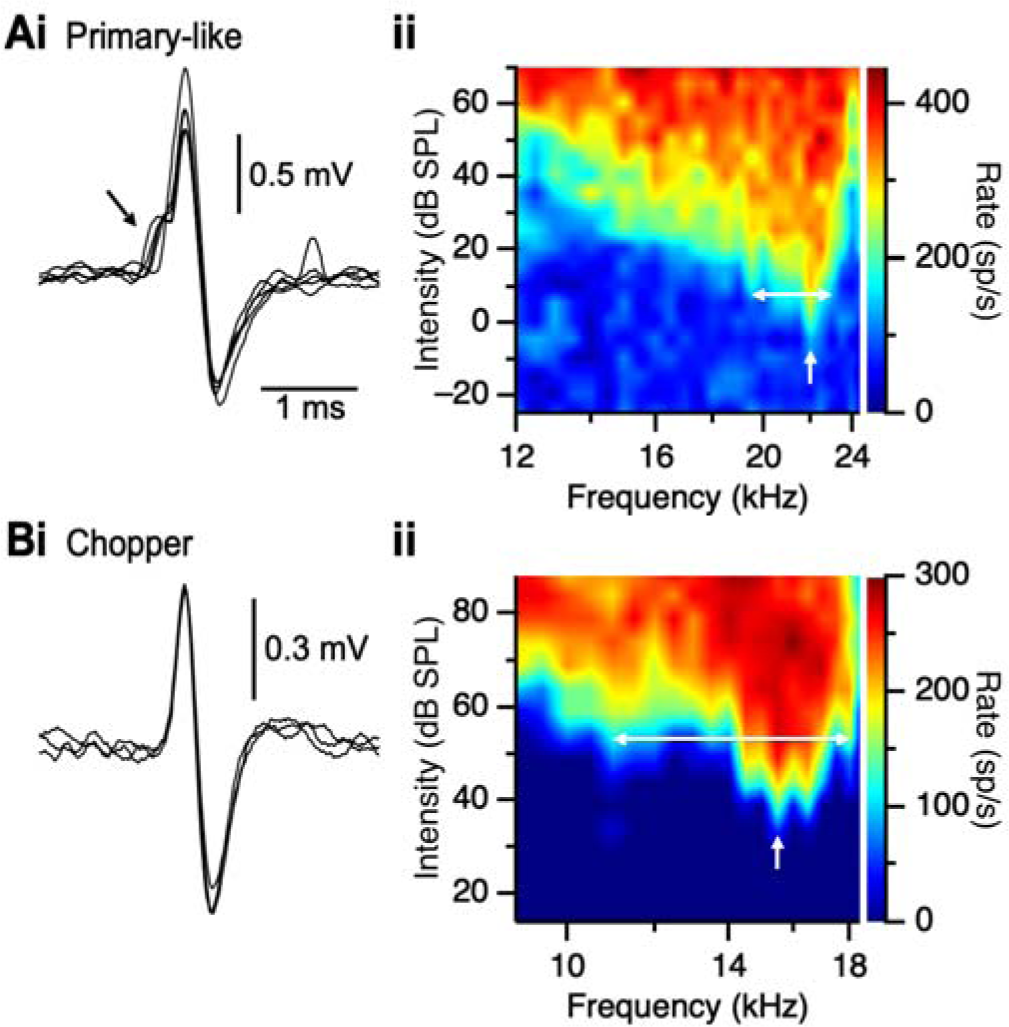
Characterizing units in the AVCN. Left panels show representative action potential waveforms for a primary-like unit with a preceding smaller EPSP (arrow, **Ai**) and a chopper with a simple spike (**Bi**). Right panels show tuning curves for units from 3–6 mo. (**Aii**) and 1.5 y CBA/CaJ (**Bii**) mice. White arrows indicate the CF of the unit, and white, horizontal lines indicate the bandwidth of *Q*_10dB_.

To evaluate how responses of AVCN units change with age, we compared responses to tones at each unit’s CF with and without masking noise. To facilitate comparisons between units, we used masking noise at a constant sensation level (20 or 30 dB above noise threshold) across units. We first quantified each unit’s sensitivity to masking noise by delivering 40 ms noise bursts at 5 dB intervals and measuring the spike rate. Representative rate-level functions (RLFs) are shown in Figure 2Ai for a primary-like unit from a 4 mo. C57 mouse, and Figure 2Bi for a chopper unit from an 8 mo. C57 mouse. We identified the noise threshold as the level at which the unit’s firing rate was significantly elevated above baseline (arrows in Fig. 2Ai, Bi) and set subsequent masking noise for each unit to 20 or 30 dB above threshold.

**Figure 2.**
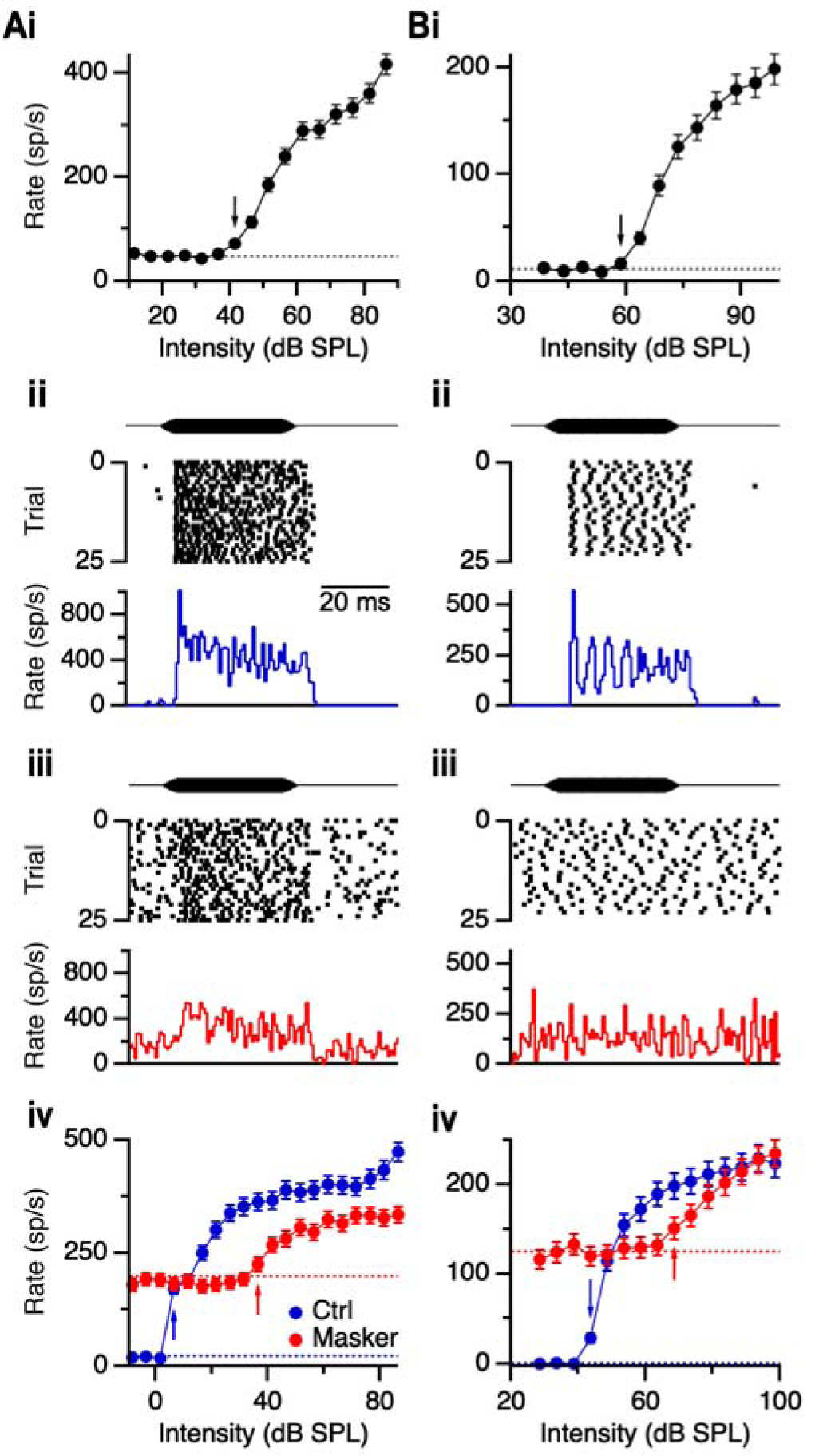
Effects of masking noise on responses to tones. (**A, B**) Representative responses of a primary-like unit (4 mo. C57, CF 17.5 kHz) (**A**) and a chopper (8 mo. C57, CF 4 kHz) (**B**). Top panels (**Ai**, **Bi**) show RLFs for 40-ms noise bursts. Dashed lines indicate spontaneous firing rate (45.5 sp/s in **A**, 11.4 sp/s in **B**), and black arrows indicate noise levels just above threshold. Middle panels show rasters and PSTHs in response to 40 ms CF tones (primary-like: 17.5 kHz, 56.5 dB SPL; chopper: 4 kHz, 58.6 dB SPL) without (**Aii**, **Bii**) and with masking noise at 30 dB above noise threshold (71.5 and 88.6 dB SPL in **Aiii** and **Biii**, respectively). Bottom panels show RLFs for the primary-like (**Aiv**) and chopper (**Biv**) units without (blue) and with (red) masking noise, with arrows indicating threshold. Dashed lines show the spontaneous firing rate without (blue) and with (red) the noise masker (22.0 and 198.1 sp/s in **Aiv**, 0.1 and 125.4 sp/s in **Biv**, respectively).

Next, we measured the responses of units to CF tones in the absence (Fig. 2Aii, 2Bii) and presence (Fig. 2Aiii, 2Biii) of the noise masker. When a noise masker was presented, baseline firing rates increased substantially, as expected. However, responses of units to tones were attenuated (Fig. 2Aiii) or absent (Fig. 2Biii) in the presence of the noise masker. We generated RLFs for each unit with and without masking noise (Fig. 2Aiv, 2Biv) and measured the thresholds to tones (blue and red arrows in Fig. 2Aiv, 2Biv). In the representative examples, threshold shifted by 30 dB for the primary-like unit and 25 dB for the chopper unit. The threshold shifts are similar to those reported in cats and gerbils (Costalupes et al., 1984; Gibson et al., 1985; Huet et al., 2018).

We examined how thresholds to CF tones change in aging CBA/CaJ mice. In primary-like units, thresholds were significantly elevated in 1.5 y CBA/CaJ mice (37.7 ± 20.8 dB SPL, 25 cells) compared to 3–6 mo. mice (10.9 ± 7.5 dB SPL, 21 cells, *p* = 0.002, K-W test) and 1 y mice (15.4 ± 5.8 dB SPL, 22 cells, *p* = 0.008, K-W test) (Fig. 3A). Thresholds of units from 1 y mice did not differ from 3–6 mo. mice (*p* > 0.05; K-W test). Furthermore, all age groups showed significant threshold shift in the presence of the noise masker (*p* < 0.001 each group, Wilcoxon signed rank test, Fig. 3B). Surprisingly, the threshold shifted significantly less for primary-like units from 1.5 y CBA/CaJ mice (25 ± 15 dB, 25 cells) compared to 3–6 mo. (40 ± 10 dB, 21 cells, *p* = 0.002, K-W test) and 1 y mice (35 ± 5 dB, 21 cells, *p* = 0.027, K-W test).

**Figure 3.**
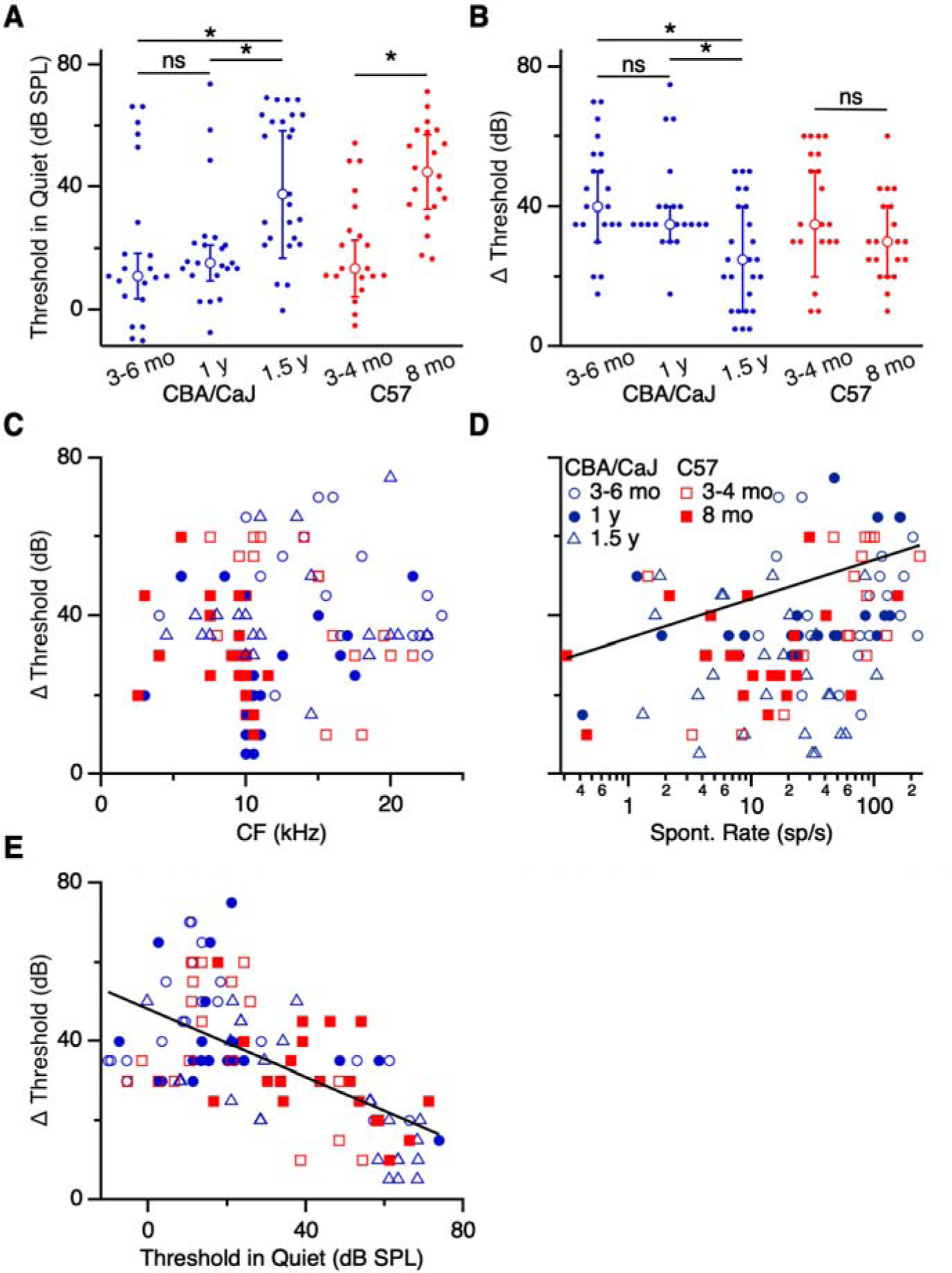
Effects of masking noise on threshold in primary-like units. (**A**) Thresholds in quiet conditions for primary-like units in CBA/CaJ (blue) and C57 (red) of different age groups. Thresholds of units from 1.5 y CBA/CaJ were significantly elevated compared to 3–6 mo. and 1 y mice (*p* < 0.01). Thresholds of units from 8 mo. C57 mice were significantly elevated compared to 3–4 mo. C57 mice (*p* < 0.001). (**B**) Shifts in threshold with the addition of masking noise. Threshold shift was significantly smaller in units from 1.5 y CBA/CaJ mice compared to 3–6 mo. and 1 y (*p* < 0.03). No age-related difference was observed for C57 mice (*p* > 0.05). (**C-E**) Threshold shift of units as a function of CF (**C**), spontaneous rate (**D**), and threshold in quiet (**E**). Threshold shift was significantly associated with spontaneous rate and threshold in quiet (*p* < 0.001). Asterisks indicate significant differences (*p* < 0.05). Summary statistics are shown throughout as median ± median absolute deviation.

To test if the change in threshold shift might be related to hearing loss in the oldest age group, we performed similar recordings and analyses in C57 mice. We reasoned that if changes in responses resulted from afferent loss, as opposed to some non-specific effect of aging, they might occur at younger ages in C57 mice, which have early-onset hearing loss. Primary-like units in 8 mo. C57 mice had significantly elevated thresholds in quiet conditions (44.8 ± 12.2 dB SPL, 20 cells) compared to 3–4 mo. mice (13.5 ± 9.1 dB SPL, 20 cells, *p* < 0.001, M-W U test; Fig. 3A), consistent with early-onset hearing loss. When the noise masker was present, thresholds increased significantly for both 3–4 and 8 mo. C57 mice (*p* < 0.001, 20 cells both groups, Wilcoxon signed rank test, Fig. 3B). However, the degree of threshold shift with masking noise did not differ significantly between the C57 age groups (3–4 mo.: 35 ± 15 dB; 8 mo: 30 ± 10 dB; *p* = 0.06, M-W U test, Fig. 3B).

To understand the cause of the decreased threshold shift in primary-like units from 1.5 y CBA/CaJ mice, we first considered if it might result from some other change in our sample, such as frequency tuning. We assessed the degree of threshold shift for units of different CF (Fig. 3C), which showed that our sample included units over a wide range of CF at all ages, with the exception of high CF units from 8 mo. C57 mice, indicative of age-related hearing loss in that group. Therefore, the decrease in threshold shift observed in units from 1.5 y CBA/CaJ mice was not likely related to CF.

Another possible factor influencing threshold shift in old mice might relate to the observation that threshold shifts are larger with louder masking noise (Costalupes et al., 1984), suggesting that the effects of masking noise might depend on a unit’s sensitivity to sound. A unit’s ANF input sensitivity is associated with its spontaneous rate, where ANFs with high spontaneous rate have high sensitivity, and ANFs with low spontaneous rate have low sensitivity (Liberman, 1978; Taberner and Liberman, 2005; Huet et al., 2016; Suthakar and Liberman, 2021). We examined how input sensitivity affected primary-like responses by examining how threshold shift varied with spontaneous rate. There was a significant association (*R^2^* = 0.17, *β* = 0.12 dB/(sp/s), *p* < 0.001; Fig. 3D), where units with low spontaneous rates had smaller shifts in threshold compared to units with high spontaneous rates. We also examined the relationship between threshold shift and threshold in quiet, which also showed a significant association (*R^2^* = 0.38, *β* = -0.43 dB/(dB SPL), *p* < 0.001; Fig. 3E). Units with low initial thresholds showed larger shifts in threshold with masking noise compared to units with high initial thresholds. Thus, the degree of threshold shift appears to be related to sensitivity. This result suggests that the lower threshold shift in the presence of masking noise for primary-like units in 1.5 y mice (Fig. 3B) is related to the increase in their thresholds in quiet (Fig. 3A).

We performed the same analyses of thresholds in chopper units of CBA/CaJ mice and found results similar to primary-like units. Thresholds in quiet for AVCN chopper units of 1.5 y mice (35.4 ± 14.0 dB SPL, 22 cells) were significantly elevated compared to 3–6 mo. mice (20.1 ± 9.2 dB SPL, 27 cells, *p* = 0.03, K-W test) and 1 y mice (13.5 ± 5.9 dB SPL, 25 cells, *p* = 0.005, K-W test; Fig. 4A). No difference was detected between thresholds of 3–6 mo. and 1 y CBA/CaJ mice (*p* > 0.05; K-W test). When masking noise was added, thresholds for units shifted for all age groups (3–6 mo.: 35 ± 5 dB, 26 cells; 1 y: 35 ± 10 dB, 25 cells; 1.5 y: 35 ± 5 dB, 22 cells; *p* < 0.001 all groups, Wilcoxon signed rank test, Fig. 4B). However, there was no significant difference in threshold shift between ages (*p* = 0.57; K-W test). In C57 mice, choppers had higher thresholds in 8 mo. mice (55.1 ± 8.7 dB SPL, 20 cells) than in 3–4 mo. mice (23.5 ± 13.2 dB SPL, 25 cells, *p* < 0.001, M-W U test; Fig. 4A). However, significant differences in threshold shifts were not observed between the two age groups (3–4 mo.: 32.5 ± 2.5 dB, 24 cells; 8 mo.: 25 ± 10 dB, 20 cells; *p* = 0.51, M-W U test; Fig. 4B).

**Figure 4.**
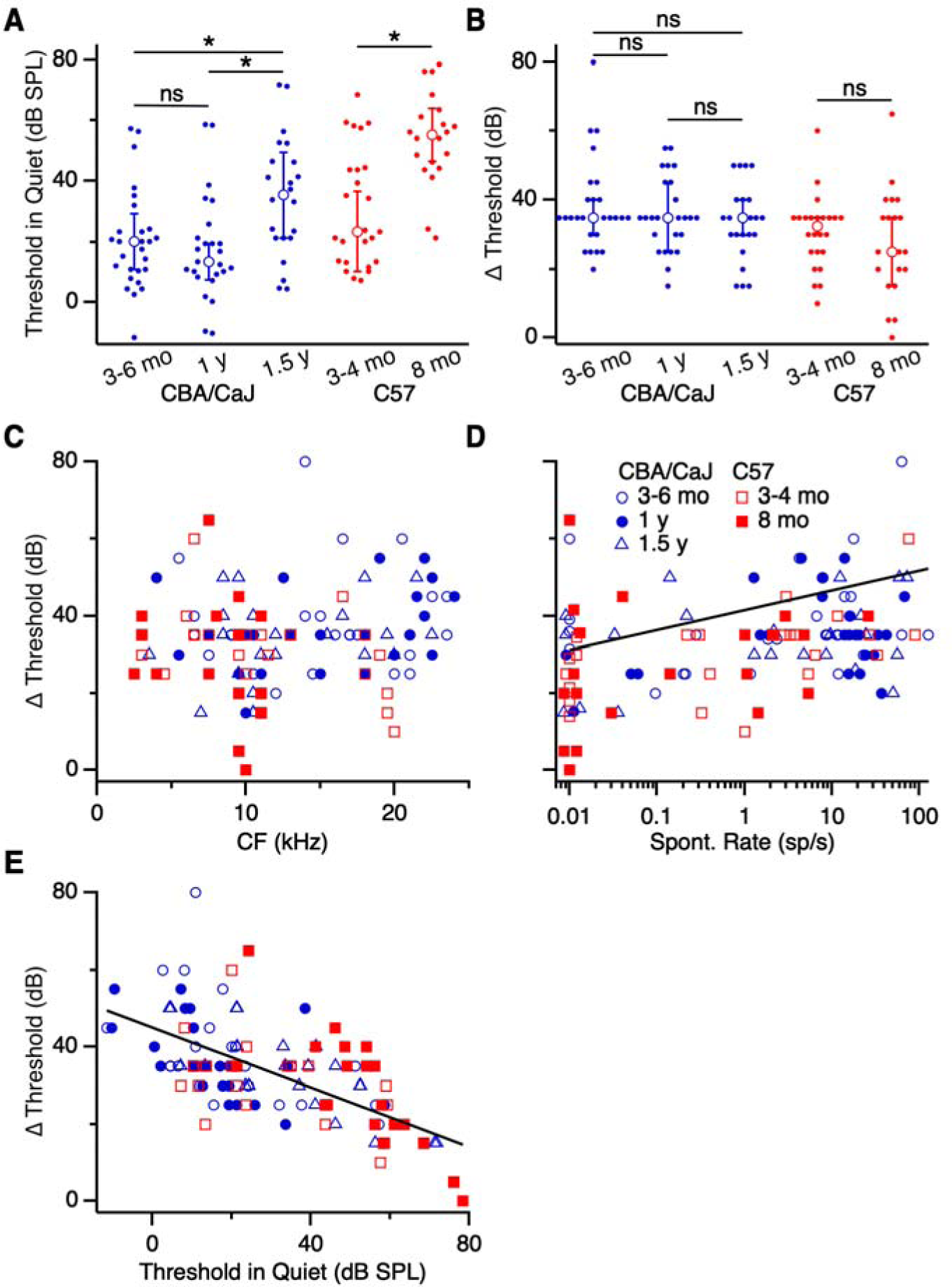
Effects of masking noise on threshold in choppers. (**A**) Thresholds in quiet conditions for primary-like units in CBA/CaJ (blue) and C57 (red) of different age groups. Thresholds of units from 1.5 y CBA/CaJ mice were significantly elevated compared to 3–6 mo. and 1 y old mice (*p* < 0.04). Thresholds of units from 8 mo. C57 mice were elevated compared to 3–4 mo. mice (*p* < 0.001). (**B**) Threshold shifts in masking noise. There were no significant differences in threshold shift between age groups (*p* > 0.05). (**C-E**) Threshold shift as a function of CF (**C**), spontaneous rate (**D**), and threshold in quiet (**E**). Threshold shift was significantly associated with spontaneous rate and threshold in quiet (*p* < 0.002).

Similar to what we saw with primary-like units, threshold shifts in choppers did not appear to depend systematically on CF (Fig. 4C). We encountered few chopper units in 8 mo. C57 mice that had CF above 11 kHz (Fig. 4C). Threshold shift did show a significant association with spontaneous rate (*R^2^* = 0.28, *β* = 0.17 dB/(sp/s), *p* < 0.002; Fig. 4D), as we observed with primary-like units (Fig. 3D). Choppers with low spontaneous rate showed smaller shifts in threshold compared to choppers with high spontaneous rate. In addition, there was a significant negative association between threshold shift and threshold in quiet (*R^2^* = 0.42, *β* = –0.39 dB/(dB SPL), *p* < 0.001; Fig. 4E). Choppers with low thresholds had greater threshold shifts than choppers with high thresholds. Thus, the sensitivity of chopper units appears to predict the degree of their threshold shift, similar to primary-like units.

### Spontaneous activity

Spontaneous activity in ANFs and the cochlear nucleus can change with age and noise exposure (Brozoski et al., 2002; Finlayson and Kaltenbach, 2009; Vogler et al., 2011; Heeringa et al., 2020; Heeringa et al., 2023). It has been suggested that high spontaneous activity in older mice is associated with decreased signal-to-noise ratio, which can impair detection of salient sound stimuli (Willott, 1990). It is possible that masking noise could exacerbate these detection impairments by elevating the baseline firing rate even further. We assessed how spontaneous activity of AVCN units changed with age in our recordings. First, considering primary-like units, the spontaneous rate decreased significantly in 1.5 y CBA/CaJ mice (18.3 ± 14.5 sp/s, 25 cells) compared to 3–6 mo. mice (78.3 ± 49.7 sp/s, 21 cells, *p* < 0.001, K-W test; Fig. 5A). No significant change in spontaneous rate was found between 3–6 mo. and 1 y CBA/CaJ mice (29.2 ± 22.0 sp/s, 22 cells, *p* > 0.05, K-W test; Fig. 5A).

**Figure 5.**
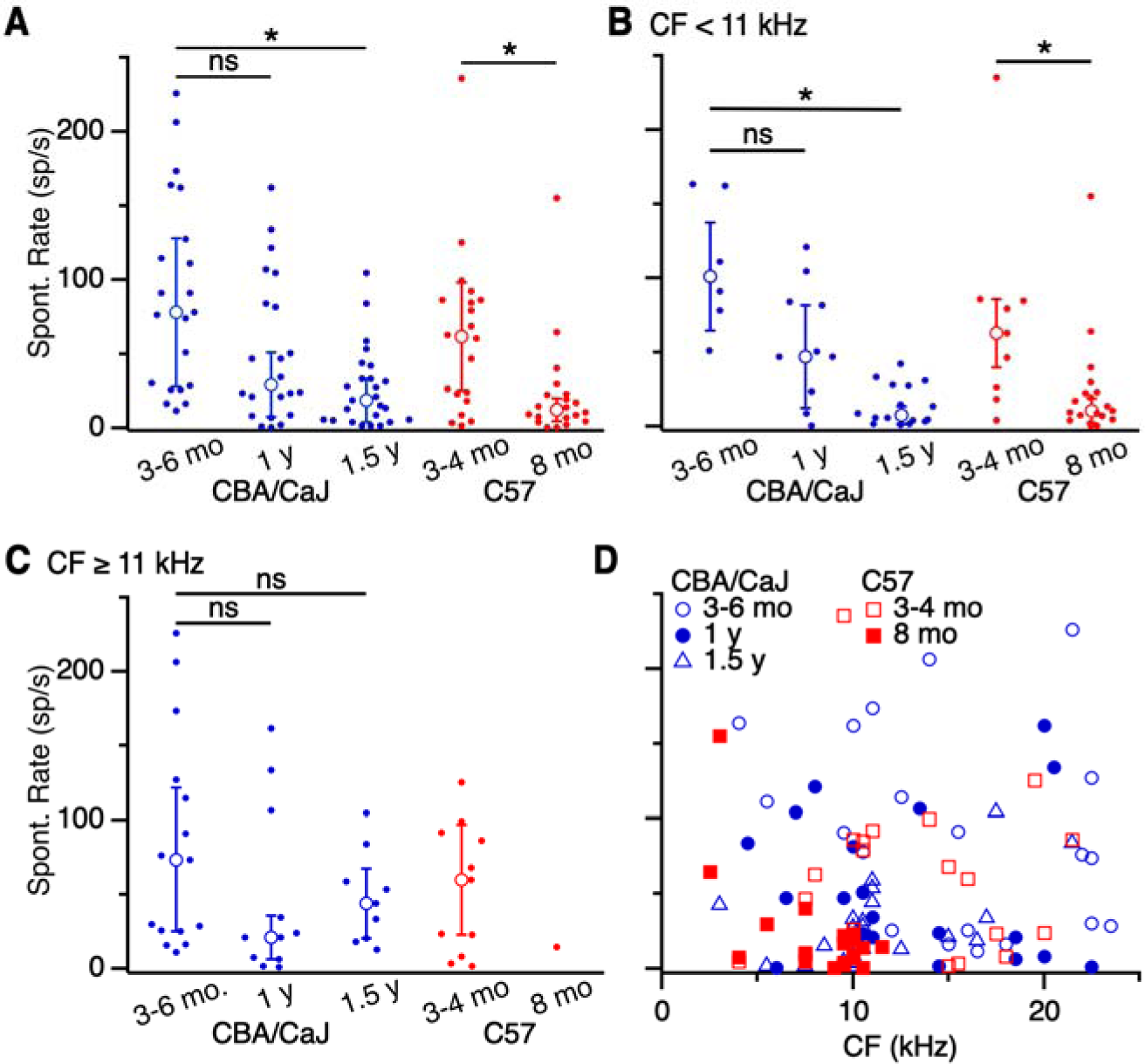
Spontaneous rate of primary-like units. (**A–C**), Spontaneous rate of (**A**) primary-like units of all CFs, (**B**) units with CF < 11 kHz, and (**C**) units with CF ≥ 11 kHz. Spontaneous rate decreased significantly with age for all primary-like units from 1.5 y CBA/CaJ and 8 mo. C57 mice (*p* < 0.006). The decrease was significant for units with (**B**) CF < 11 kHz (*p* < 0.01), but (**C**) not CF ≥ 11 kHz (*p* > 0.05). (**D**) Spontaneous rates for units of different CFs. There was no significant relationship between CF and spontaneous rate (*p* > 0.05). Asterisks indicate significant differences (*p* < 0.05).

We wondered if high-frequency units might show differential effects, related to high-frequency hearing loss. We found fewer units with CF > 11 kHz in 8 mo. C57 mice, suggesting these are particularly sensitive during the onset of hearing loss, so we categorized units into CF < 11 kHz or ≥ 11 kHz. Spontaneous rate of primary-like units with a CF < 11 kHz decreased significantly for 1.5 y CBA/CaJ mice (7.3 ± 5.8 sp/s, 16 cells) compared to 3–6 mo. mice (101.1 ± 36.3 sp/s, 6 cells, *p* < 0.001, K-W test), but no difference was detected at 1 y (47.0 ± 34.5 sp/s, 12 cells; *p* = 0.11, K-W test; Fig. 5B). By contrast, we did not observe age-related changes in spontaneous rate at CF ≥ 11 kHz for CBA/CaJ mice (3–6 mo.: 73.7 ± 48.2 sp/s, 15 cells; 1 y: 21.1 ± 14.6 sp/s, 11 cells; 1.5 y: 44.0 ± 23.3 sp/s, 9 cells; *p* > 0.23, K-W test; Fig. 5C).

We performed similar analyses on primary-like units in C57 mice. Spontaneous rates decreased significantly in C57 mice for all units grouped together (3–4 mo.: 61.5 ± 36.5 sp/s, 20 cells; 8 mo.: 12.0 ± 7.6 sp/s, 20 cells, Fig. 5A) and units with CF < 11 kHz (3–4 mo.: 62.9 ± 23.2 sp/s, 9 cells; 8 mo.: 10.3 ± 8.1 sp/s, 19 cells, Fig. 5B) (*p* < 0.01 both comparisons, M-W U test). For CF ≥ 11 kHz, our sample only included a single primary-like unit in 8 mo. C57, which precluded a meaningful analysis of the effects of age.

To determine whether our recordings were reasonably representative of the frequency diversity of units, we plotted spontaneous rate of units against their CF (Fig. 5D). It appeared that our sample included varied spontaneous rates across CF for primary-like units, except there were few high-CF units in 8 mo. C57 mice, consistent with the well-known early-onset high frequency hearing loss in this strain (Xiong et al., 2020).

We performed similar analyses on chopper units. In units from CBA/CaJ mice, we found no age-related changes in spontaneous rate (3–6 mo.: 6.7 ± 6.6 sp/s, 27 cells; 1 y: 13.9 ± 10.4 sp/s, 25 cells; 1.5 y: 3.5 ± 3.5 sp/s, 22 cells; *p* > 0.05, K-W test; Fig. 6A). No significant age-related changes were observed in spontaneous rate in units even after subdividing for CF < 11 kHz (3–6 mo.: 6.7 ± 4.8 sp/s, 9 cells; 1 y: 13.9 ± 10.4 sp/s, 7 cells; 1.5 y: 0.2 ± 0.2 sp/s, 11 cells) or CF ≥ 11 kHz (3–6 mo.: 6.9 ± 6.9 sp/s, 18 cells; 1 y: 14.9 ± 11.7 sp/s, 18 cells; 1.5 y: 8.8 ± 8.8 sp/s, 11 cells) (both *p* > 0.4, K-W tests, Fig. 6B,C). In C57 mice, we also did not find a change in spontaneous rate with age when all chopper units were evaluated (3–4 mo.: 1 ± 1 sp/s, 25 cells; 8 mo.: 0.035 ± 0.035 sp/s, 20 cells; *p* = 0.12, M-W U Test; Fig. 6A) and for units with CF ≥ 11 kHz (3–4 mo.: 0.7 ± 0.7 sp/s, 14 cells; 8 mo.: 2.2 ± 1.3 sp/s, 4 cells; *p* = 0.57, M-W U test; Fig. 6C). However, when spontaneous rate was examined in chopper units with CF < 11 kHz, a significant decrease was found in units from 8 mo. C57 mice (0 ± 0 sp/s, 16 cells) compared to 3–4 mo. mice (2.8 ± 2.8 sp/s, 11 cells) (*p* = 0.03, M-W U test; Fig. 6B). Our sample of choppers also showed varied spontaneous rate across all CFs (Fig. 6D), with the exception of spontaneous rates of units from 8 mo. C57 mice that were clustered in the lower frequency range. These results suggest that changes in spontaneous rate are relatively minor in chopper units.

**Figure 6.**
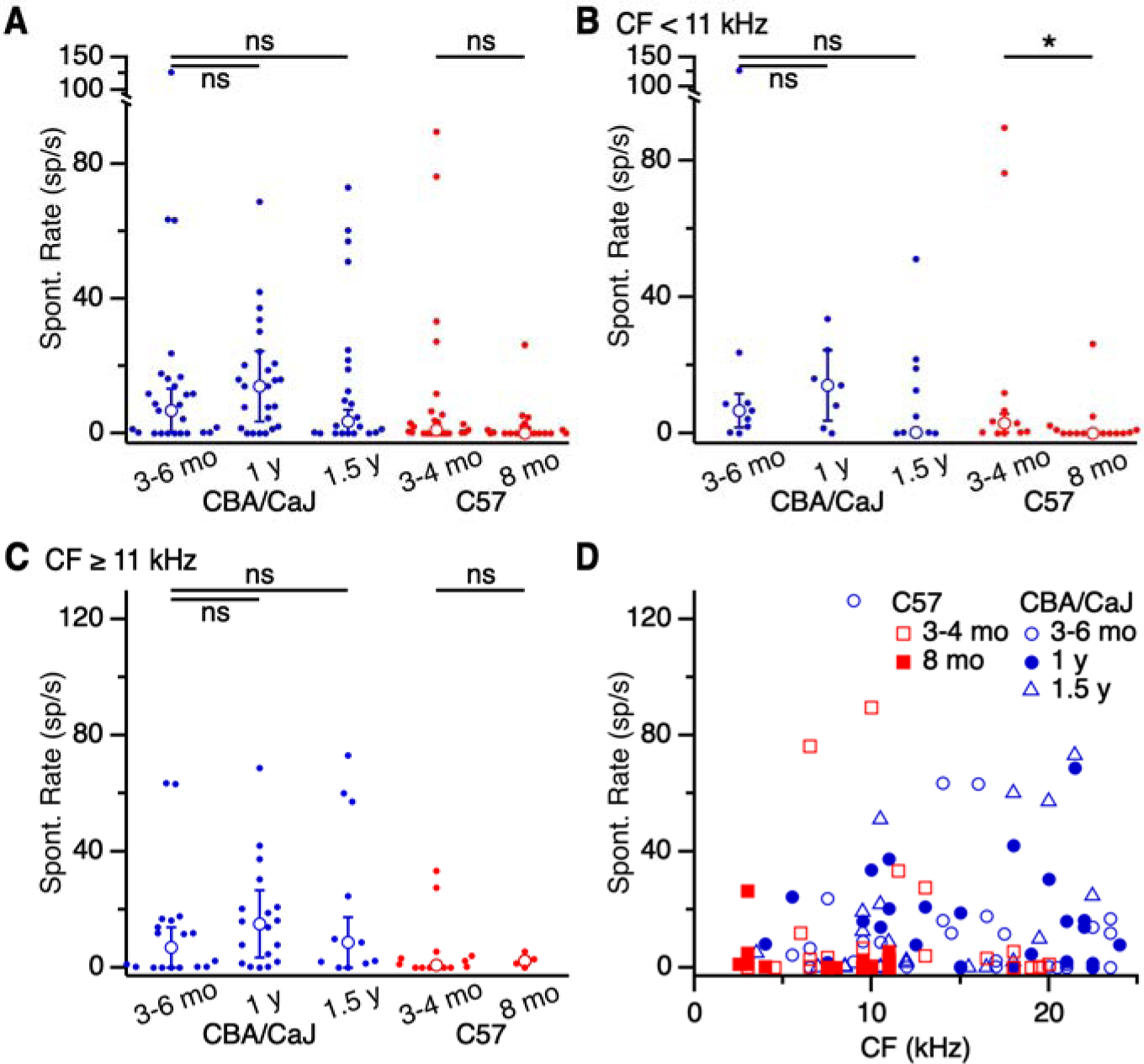
Changes in spontaneous rate of choppers with age. (**A–C**) Spontaneous rate of (**A**) choppers of all CFs, (**B**) choppers with CF < 11 kHz, and (**C**) choppers with CF ≥ 11 kHz. There were no significant changes with age, except choppers from 8 mo. C57 mice (*p* = 0.03). (**D**) Spontaneous rates for choppers of different CFs. There was no significant relationship between CF and spontaneous rate (*p* > 0.05).

Overall, spontaneous activity appeared to decrease with age in primary-like and chopper units, particularly for units with CF < 11 kHz. This is consistent with the observation that spontaneous rate decreases with age in ANF units with low CFs (Heeringa et al., 2020).

### Dynamic range

Dynamic range can narrow in auditory units after moderate and traumatic noise exposure (Gao et al., 2009; Bures et al., 2010; Cheng et al., 2017), which could lead to impaired detection and perception of salient sound stimuli, particularly in the presence of masking noise. We therefore examined how dynamic range changes with aging. Figure 7A shows a representative RLF from a chopper unit in a 5 mo. CBA/CaJ mouse, fit to a sigmoid curve. We quantified the dynamic range as the difference in tone intensities associated with firing rates at 10% and 90% of the maximum driven rate (dotted lines in Fig. 7A), which was 43 dB for this unit. We analyzed units with clearly-saturating RLFs (36–64% of each group’s total), and found no significant age-related change in dynamic range across all strains and ages for primary-like units (CBA/CaJ 3–6 mo.: 24.7 ± 4.4 dB, 13 cells; 1 y: 26.9 ± 8.9 dB, 14 cells; 1.5 y: 24.6 ± 4.4 dB, 10 cells; *p* = 0.93, K-W test; C57 3–4 mo.: 18.2 ± 6.4 dB, 12 cells; 8 mo.: 23.8 ± 6.3 dB, 10 cells; *p* = 0.77, M-W U test; Fig. 7B) or choppers (CBA/CaJ 3–6 mo.: 20.3 ± 7.8 dB, 14 cells; 1 y: 17.2 ± 7.5 dB, 16 cells; 1.5 y: 17.4 ± 7.4 dB, 11 cells; *p* = 0.39, K-W test; C57 3–4 mo.: 12.2 ± 10.8 dB, 9 cells; 8 mo.: 19.2 ± 3.3 dB, 9 cells; *p* = 0.44, M-W U test; Fig. 7D).

**Figure 7.**
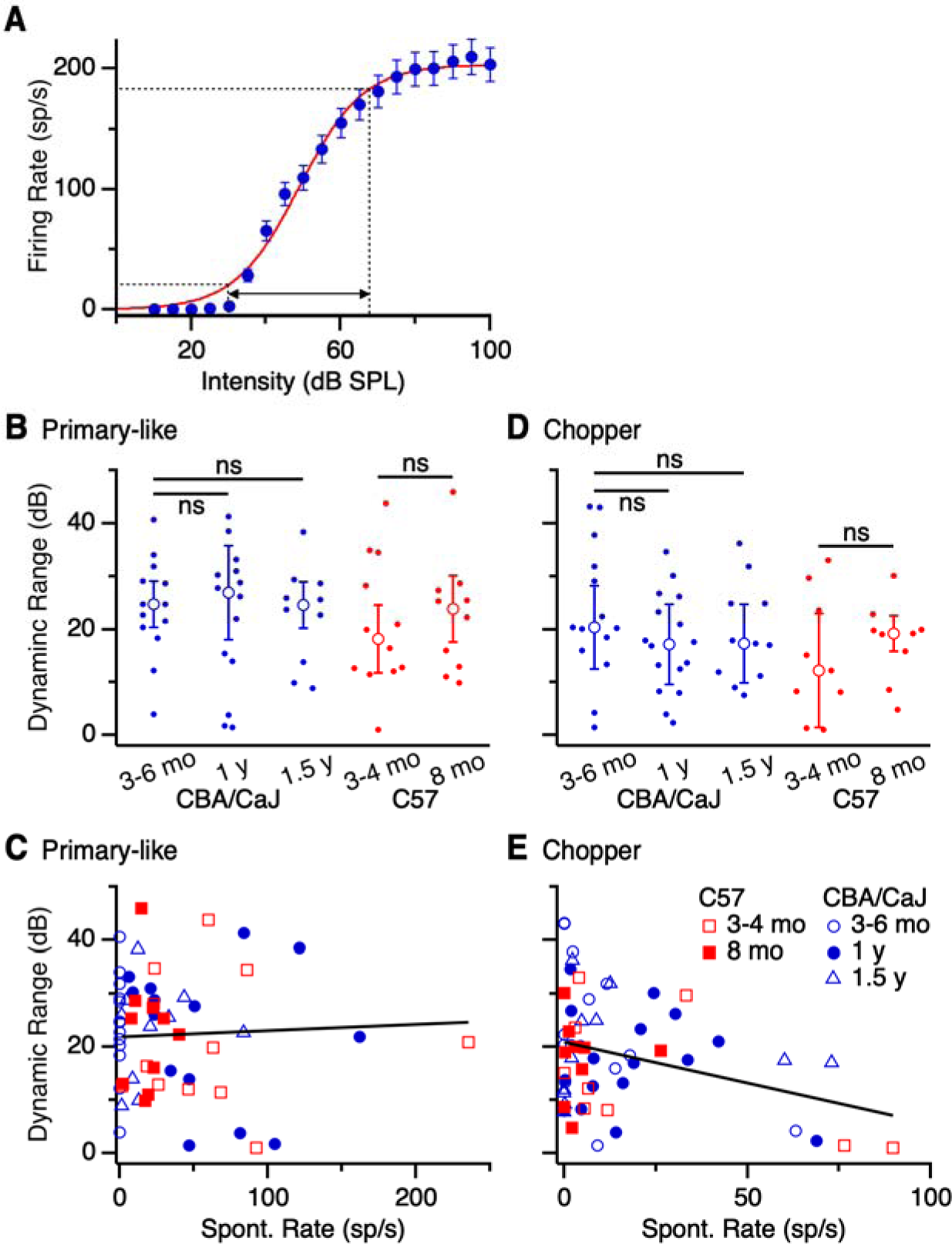
Effects of age on dynamic range of primary-like units and choppers. (**A**) Representative dynamic range of a chopper from a 5 mo. CBA/CaJ mouse. The RLF was fit to a sigmoid function. Horizontal dotted lines indicate 10 and 90% of the maximum spike rate, and vertical lines indicate the dynamic range. (**B, D**) Dynamic range for (**B**) primary-like units and (**D**) choppers in different age categories. No significant changes were observed with age (*p* > 0.05). (**C, E**) Dynamic range for (**C**) primary-like units and (**E**) choppers as a function of spontaneous rate. There was a significant negative association between dynamic range and spontaneous rate for chopper units (*p* = 0.012), but no significant relationship for primary-like units (*p* > 0.05).

Previous work has demonstrated that dynamic range is largest in low spontaneous rate ANFs (Evans and Palmer, 1980; Taberner and Liberman, 2005; Huet et al., 2016). To test if this relationship holds for aging AVCN principal cells, we examined the relationship between dynamic range and spontaneous rate in our sample of AVCN units. There was no significant relationship in primary-like units (*R^2^* = 0.003, *β* = 0.011 dB/(sp/s), *p* = 0.7; Fig. 7C), but there was a significant negative association in chopper units (*R^2^* = 0.13, *β* = –0.17 dB/(sp/s), *p* = 0.012; Fig. 7E). This suggests that choppers with lower spontaneous rates had wider dynamic ranges than choppers with higher spontaneous rates, similar to ANFs (Taberner and Liberman, 2005).

### Tuning precision

Previous work has reported broader frequency tuning in ANFs from noise-exposed and aged mice (Ohlemiller, 2002). Broader frequency tuning could reduce frequency discrimination, and affect detection of tones in the presence of masking noise. To evaluate how tuning precision changes with age in our dataset, we measured the *Q*_10dB_ of units by taking the ratio of CF to the bandwidth at 10 dB above threshold (examples in Fig. 1). We detected no age-related changes in CBA/CaJ or C57 mice in *Q*_10dB_ of primary-like units (CBA/CaJ 3–6 mo.: 5.6 ± 2.0, 13 cells; 1 y: 5.1 ± 0.9, 14 cells; 1.5 y: 4.4 ± 1.5, 13 cells; *p* = 0.77, K-W test; C57 3–4 mo.: 4.2 ± 0.5, 9 cells; 8 mo.: 2.8 ± 0.9, 16 cells, *p* = 0.07, M-W U test; Fig. 8A) or chopper units (CBA/CaJ 3–6 mo.: 5.5 ± 1.5, 22 cells; 1 y: 6.2 ± 1.0, 18 cells; 1.5 y: 3.5 ± 1.3, 11 cells; *p* = 0.06, K-W test; C57 3–4 mo.: 4.3 ± 1.2, 23 cells; 8 mo.: 3.1 ± 1.2, 17 cells, *p* = 0.22 M-W U test; Fig. 8B). This suggests that for units that continue to respond into old age, cochlear inputs giving rise to their responses is also largely intact. This is consistent with previous findings showing no change in *Q*_10dB_ values in aging gerbils (Heeringa et al., 2020).

**Figure 8.**
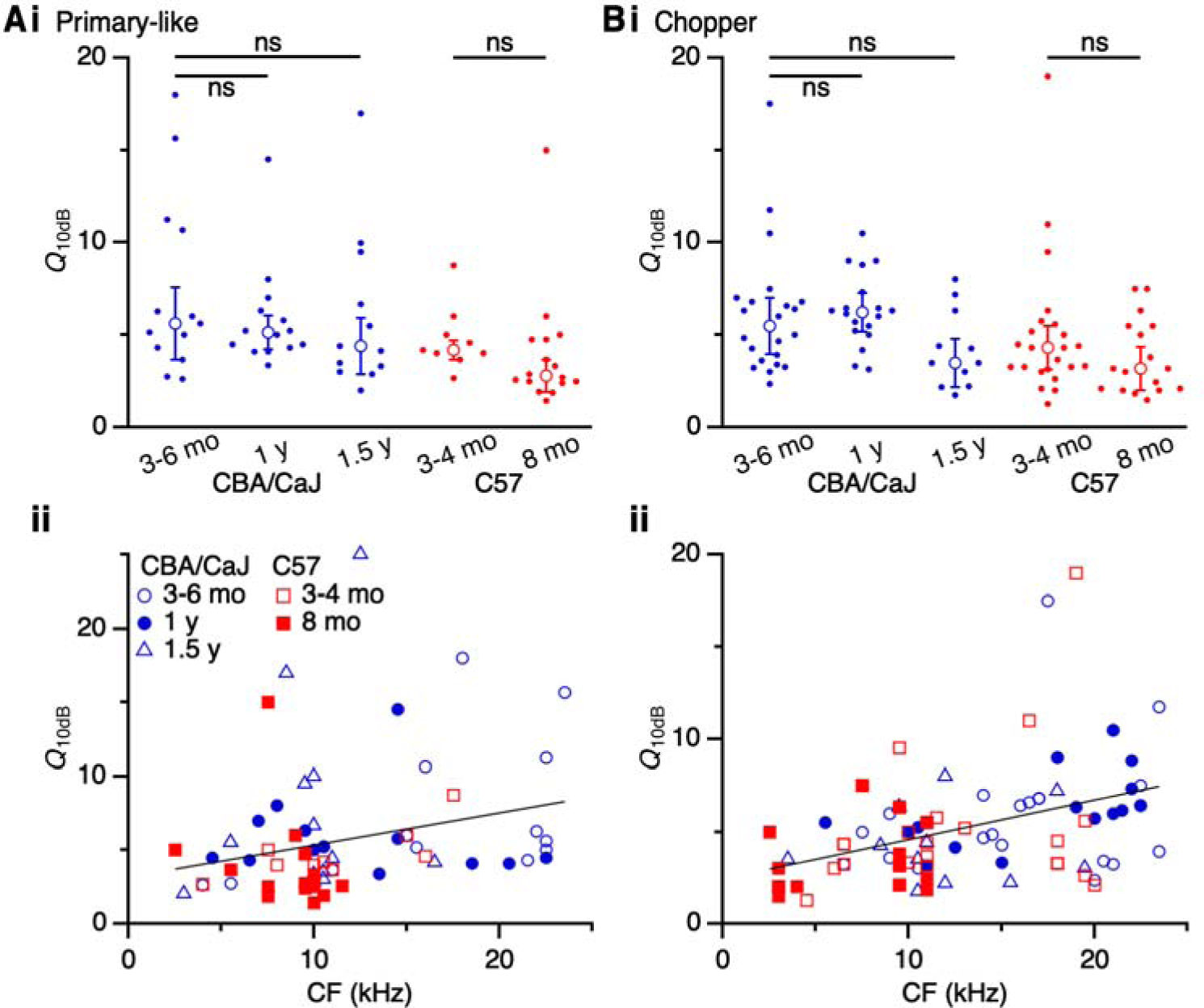
Effects of age on tuning precision in primary-like and chopper units. *Q*_10dB_ for (**A**) primary-like units and (**B**) choppers in CBA/CaJ and C57 mice. Upper panels, *Q*_10dB_ across age groups, showing no significant changes (*p* > 0.05). Lower panels, *Q*_10dB_ across CF in primary-like units (**Aii**) and choppers (**Bii**), which both showed significant positive associations (*p* < 0.033).

We assessed the relationship between sharpness of tuning and CF, which has been previously demonstrated in CBA/CaJ and C57 mice (Taberner and Liberman, 2005). We found significant positive associations for primary-like units (*R^2^* = 0.071, *β* = 0.22 kHz^-1^, *p* = 0.032; Fig. 8Aii) and choppers (*R^2^* = 0.17, *β* = 0.21 kHz^-1^, *p* < 0.001; Fig. 8Bii). Units with the lowest CFs had low *Q*_10dB_ values, while units with the highest CFs had high *Q*_10dB_ values.

### Spike timing

Finally, neurons of the AVCN play a key role in encoding the temporal features of sounds, for such functions as sound localization. We evaluated changes in temporal precision of AVCN units by quantifying the first spike latency (FSL) and jitter for CF tones 10 dB above threshold. Primary-like units showed a slight increase in FSL with age in CBA/CaJ mice (3–6 mo.: 7.1 ± 1.0 ms, 19 cells; 1 yr: 7.8 ± 1.5 ms, 21 cells; 1.5 yr: 8.5 ± 1.6 ms, 24 cells; *p* = 0.042, K-W test; posthoc Dunn’s test, 3–6 mo. vs. 1.5 yr *p* = 0.035, other comparisons *p* > 0.35) and C57 mice (3–4 mo.: 7.3 ± 1.0 ms; 8 mo.: 9.5 ± 1.6 ms; *p* = 0.003, M-W U test) (Fig. 9A). There was no significant change in jitter in either strain with age (CBA/CaJ 3–6 mo.: 1.0 ± 0.3 ms, 19 cells; 1 yr: 1.4 ± 0.6 ms, 21 cells; 1.5 yr: 1.5 ± 0.6 ms, 24 cells; *p* = 0.07 KW test; C57 3–4 mo.: 1.6 ± 0.9 ms, 20 cells; 8 mo.: 1.4 ± 0.5 ms, 19 cells; *p* = 0.58, M-W U test) (Fig. 9B). Thus, while primary-like response latencies were slightly delayed with age, the precision of those spikes appeared to remain consistent across ages.

**Figure 9.**
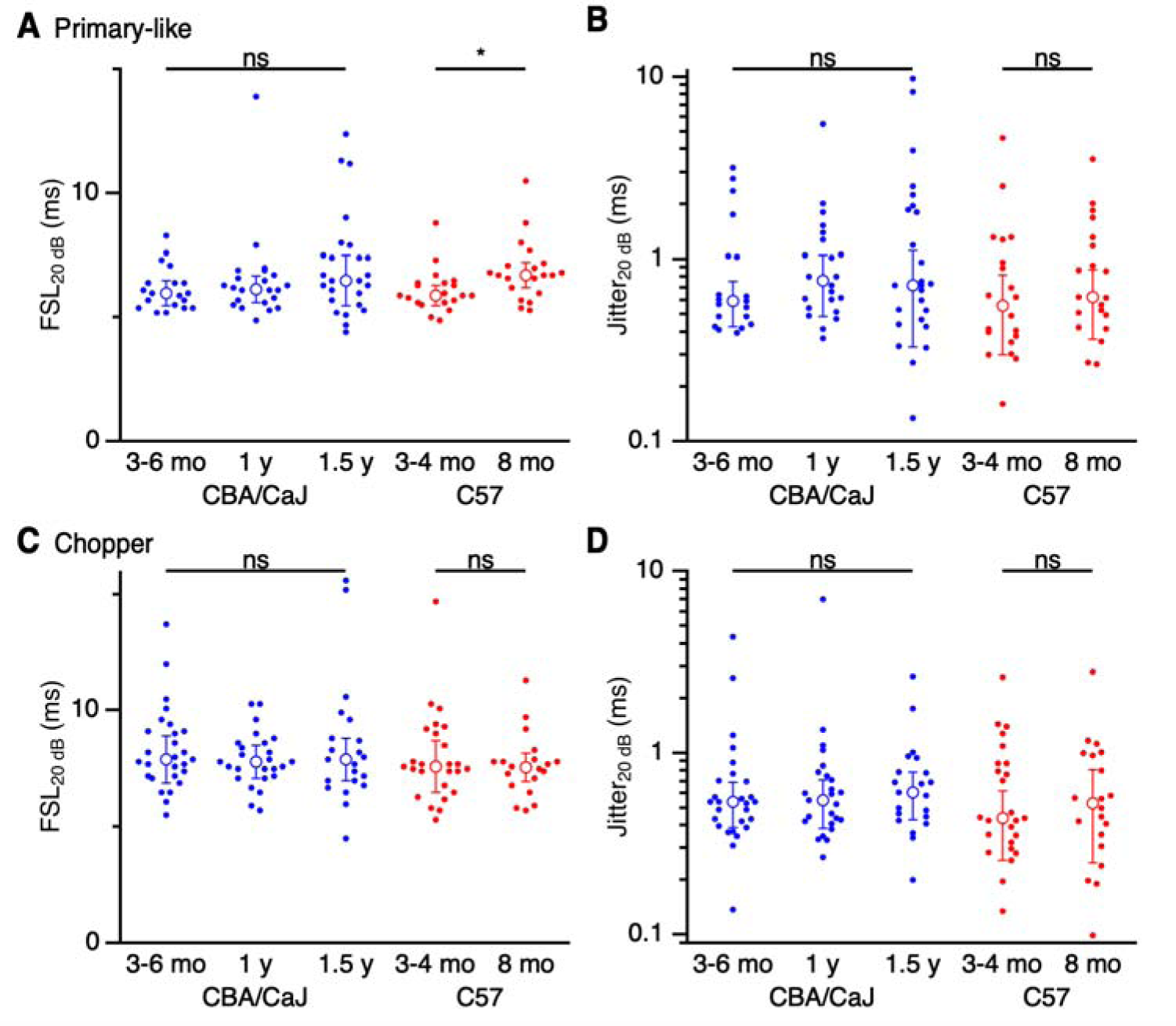
Effects of age on first spike latency (FSL) and jitter, measured at 10 dB above threshold. (**A**) FSL_10 dB_ for primary-like units in CBA/CaJ (blue) and C57 (red) mice. Latency increased significantly with age in both groups (*p* < 0.05). (**B**) Jitter_10 dB_ for primary-like units in CBA/CaJ (blue) and C57 (red) mice. Jitter showed no significant changes with age (*p* > 0.05). (**C**, **D**) Choppers showed no significant change in latency (**C**) or jitter (**D**) in older CBA/CaJ or C57 mice (*p* > 0.5).

Choppers do not appear to encode temporal features of sounds with great precision. Furthermore, they showed no significant age-related latency changes for either mouse strain (CBA/CaJ 3–6 mo.: 10.1 ± 1.8 ms, 27 cells; 1 yr: 10.6 ± 1.3 ms, 25 cells; 1.5 yr: 10.7 ± 2.5 ms, 21 cells; *p* = 0.75 KW test; C57 3–4 mo.: 9.5 ± 1.9 ms, 25 cells; 8 mo.: 9.9 ± 1.1 ms, 20 cells; *p* = 0.62, M-W U test; Fig. 9C) or jitter (CBA/CaJ 3–6 mo.: 1.1 ± 0.6 ms, 27 cells; 1 yr: 1.2 ± 0.6 ms, 25 cells; 1.5 yr: 1.2 ± 0.7 ms, 21 cells; *p* = 0.64 KW test; C57 3–4 mo.: 1.2 ± 0.7 ms, 25 cells; 8 mo.: 0.93 ± 0.47 ms, 20 cells; *p* = 0.87, M-W U test; Fig. 9D). Thus, temporal information coding in chopper units does not appear to degrade with age, consistent with their limited role in processing timing cues.

## Discussion

We examined how age-related changes in AVCN principal cells might contribute to difficulty hearing in the presence of masking noise. The AVCN is one of the first stages of the central auditory pathway, so changes there could greatly influence subsequent sound processing. We evaluated if bushy and stellate cells from old mice show greater shifts in sensitivity to tones in the presence of masking noise. However, shifts in sensitivity were the same or smaller in aged mice compared to young mice. These results suggest that age-related hearing deficits in detection of tones in noise (Kobrina et al., 2020) do not result from degraded responses of individual neurons in the AVCN. Rather, AVCN neurons maintain fairly consistent representations of tones and tones in noise into old ages.

### Age-related hearing-in-noise deficits are not explained by AVCN single unit response properties

The decrease in threshold shift in old primary-like units is consistent with behavioral and auditory brainstem responses measured in aging mice (Ehret, 1979; Vicencio-Jimenez et al., 2021). A major change during aging is diminished input from the periphery, from both death of hair cells and loss of functional synapses onto ANFs. In the AVCN, old ANF synapses onto bushy cells show increased depression and reduced temporal precision *in vitro* (Xie and Manis, 2017), with the overall effect of reduced fidelity (Xie, 2016). Such changes could reduce the impact of constant masking noise, and enhance the responses of primary-like units to salient, transient stimuli.

Furthermore, our findings suggest that age-related hearing-in-noise deficits result from disrupted processing downstream of the AVCN. One possibility is the medial olivocochlear efferents (MOC), which inhibit outer hair cell function and have been implicated in promoting detection of salient sound stimuli by reducing responses to masking noise (Fuchs and Lauer, 2019; Lauer et al., 2022). Age-related loss of MOC neurons impairs sound detection in noisy environments (Vicencio-Jimenez et al., 2021). Augmenting MOC feedback by increasing the inhibitory response in outer hair cells prevents some of the anatomical and functional auditory decline observed in older mice (Boero et al., 2020). However, MOC changes would also be expected to yield changes in cochlear responses, which from our data appear to be minor and unlikely to account for increased difficulty with speech in noise.

Another possibility is that primary neurons in other parts of the cochlear nucleus could be disrupted by reduced afferent input. One major cell type that would be valuable to consider in future studies is octopus cells, which integrate activity from multiple ANFs tuned to different frequencies (Oertel et al., 2000). This integrative function makes them especially suited for encoding temporal information in broadband sounds, such as sound onset and offset (Golding et al., 1995), which contributes to the responses of downstream targets including the superior paraolivary nucleus (Gomez-Alvarez et al., 2018; Rajaram et al., 2019). It would be valuable to determine whether this integrative function also makes octopus cells vulnerable to loss of afferent input during aging.

Age-related deficits with hearing-in-noise could also result from higher-order auditory areas, rather than the cochlear nucleus. This could be through changes in circuits at later stages of the auditory pathway, such as in inferior colliculus or auditory cortex (Harris et al., 2009; Presacco et al., 2019; Shilling-Scrivo et al., 2021, 2022) or degeneration of auditory efferent pathways (Zettel et al., 2007; Radtke-Schuller et al., 2015; Liberman and Liberman, 2019; Kobrina et al., 2020; Vicencio-Jimenez et al., 2021). Another possibility is that the detection of tones in noise requires a large volume of information coming from the periphery. That is, individual AVCN units may show fairly consistent responses with aging as we found, and by implication ANFs probably do as well. However, the processing that segregates signals such as tones or speech from noise may require integration of many inputs, which becomes more difficult with aging and peripheral cell loss.

We found that the threshold shift caused by masking noise was similar across age groups in C57 mice. This was surprising, because C57 mice show significant peripheral damage at 8 mo. (Francis et al., 2003), similar to 1.5 y CBA/CaJ mice. However, the threshold shift actually decreased in the 1.5 y CBA/CaJ group. This suggests the decreased shift in 1.5 y CBA/CaJ mice relates more to their age than simply a decrease in input from the periphery. Peripheral input is also reduced in 8 mo. C57 mice, but they show no decrease in threshold shift. In other words, 1.5 y CBA/CaJ mice appear to have a mechanism that reduces the impact of masking noise on tone thresholds, that 8 mo. C57 mice lack, so they do not show a decrease in threshold shift.

Unlike primary-like units, chopper units (stellate cells) showed no significant changes in threshold shift with age. One possible explanation for this difference is that primary-like units undergo changes as they age in the complement of excitatory and inhibitory inputs they receive, to a greater extent than chopper units. Excitation to both cell types comes exclusively from ANFs, so changes in the strength of ANF synapses could underlie this difference (Ferragamo et al., 1998; Xu-Friedman and Regehr, 2005a, b; Yang and Xu-Friedman, 2008, 2009; Oertel et al., 2011). For instance, older bushy cells appear to particularly lose synapses formed by ANF subtypes with low spontaneous rates and high thresholds (Wang et al., 2021), which could reduce the overall excitation they receive and decrease the impact of masking noise. Furthermore, the ANF synapses seem to shift from predominantly somatic to dendritic (Wang et al., 2023), which could change how excitation and inhibition interact and drive spiking. Inhibitory circuits in the auditory system also change with age (Caspary et al., 2008). Accounting for the differences between primary-like and chopper units during aging will require better understanding of changes in inhibition specifically onto these cell types.

### Spontaneous activity decreases in AVCN primary-like units

Spontaneous activity of AVCN units may provide additional insights into changes in levels of excitation and inhibition. We found that spontaneous rates decreased with age in primary-like units of both CBA/CaJ and C57 mice, particularly among units with CF < 11 kHz. By contrast, spontaneous activity in choppers changed little in our sample, and spontaneous activity appears to increase in the dorsal cochlear nucleus with age (Caspary et al., 2005; Caspary et al., 2006). These differences most likely result from distinct circuitry related to the different cell types (Young and Oertel, 2004). Spontaneous firing in ANFs declines with age (Heeringa et al., 2020), which would likely reduce excitation and spontaneous firing in both primary-like and chopper units. In addition, the volume of synaptic areas in endbulbs expressing vesicular glutamate transporter 1 (VGluT1) decreases with age in the AVCN (Wang et al., 2021), suggesting bushy cells receive less excitation. Furthermore, the accumulation of cochlear damage and synaptopathy during aging would likely reduce the number of effective auditory nerve inputs to AVCN neurons, and thereby reduce their spontaneous activity (Zeng et al., 2009; Heeringa et al., 2016; Schrode et al., 2018).

Reduced spontaneous rates could also result from an increase in the strength of inhibitory inputs with age. Inhibition in the AVCN is primarily glycinergic, so increased inhibition could result from elevated glycine levels or upregulation of glycine receptors in the AVCN. However, glycine levels and post-synaptic receptor binding sites decrease in the cochlear nucleus with age and noise-induced cochlear damage (Banay-Schwartz et al., 1989b; Wang et al., 2009; Schrode et al., 2018). Thus, increased inhibition is an unlikely explanation for the age-related decrease in spontaneous activity of primary-like units we observed.

Another possible mechanism contributing to reduced spontaneous rate could be selective loss of ANFs with high spontaneous rate, leaving an over-representation of low-spontaneous-rate ANFs. However, noise-induced damage and age-related decline are thought to have the greatest effect on low-spontaneous rate, high-threshold ANFs (Schmiedt et al., 1996; McClaskey et al., 2022). In addition, anatomical evidence indicates that most bushy cells in old mice are dominated by input from ANFs that express calretinin, which are thought to have relatively high spontaneous rates (Wang et al., 2021). Thus, it seems unlikely that spontaneous rate decreases because of increased representation of low-spontaneous-rate ANFs.

Interestingly, we found only minor changes in the spontaneous rate of chopper units. Chopper units receive excitation from ANFs and inhibition from stellate interneurons and cells of the dorsal cochlear nucleus (Ferragamo et al., 1998; Brugge, 2013; Ngodup et al., 2020). Markers for both excitatory glutamatergic and inhibitory glycinergic synapses decrease with age (Banay-Schwartz et al., 1989a). The minor change in spontaneous activity suggests that excitation and inhibition decrease relatively similarly with age, which could preserve the balance of excitation and inhibition to choppers.

### Changes in dynamic range and tuning

Noise exposure can have a dramatic impact on dynamic range and tuning. Dynamic range can widen or narrow in different auditory areas following noise exposure (DCN: Brozoski et al., 2002; AC: Gao et al., 2009; IC: Cheng et al., 2017). Noise exposure can also affect tuning (Salvi et al., 1978; Kaltenbach et al., 1992; Muller and Smolders, 2005), especially broadening unit tuning following outer hair cell damage (Liberman and Dodds, 1984; Smith et al., 1987; Cai et al., 2009).

However, we observed no significant changes in dynamic range or tuning precision of primary-like and chopper units with age. This is consistent with another study showing no change in tuning sharpness in ANF units in aging gerbils (Heeringa et al., 2020). Dynamic range also changes little in the inferior colliculus of aging rats (Palombi and Caspary, 1996). Thus, the changes that take place during aging may not be equivalent to the accumulated effects of a lifetime of incremental noise traumas.

## Acknowledgments

This work was supported by National Institutes of Health grants R01 DC015508 to MAX-F, R01 DC017620 to AML, T32 DC000023 to KB, and the David M. Rubenstein Fund for Hearing Research to AML and CJCG. The authors thank James Engel, Kimberly Nguyen, and the Laboratory Animal Facility staff at the University at Buffalo for maintaining the animal colony. We thank Dr. Micheal Dent for providing mice and recording equipment. We also thank Drs. Wei Sun and Richard Salvi for providing sound calibration equipment.

